# VPS13D bridges the ER to Miro containing membranes

**DOI:** 10.1101/2020.10.07.328906

**Authors:** Andrés Guillén-Samander, Marianna Leonzino, Michael G. Hanna, Ni Tang, Hongying Shen, Pietro De Camilli

## Abstract

Mitochondria, which are excluded from the secretory pathway, depend on lipid transport proteins for their lipid supply from the ER, where most lipids are synthesized. In yeast, the outer mitochondrial membrane GTPase Gem1 is an accessory factor of ERMES, an ER-mitochondria tethering complex that contains lipid transport domains and that functions, partially redundantly with Vps13, in lipid transfer between the two organelles. In metazoa, where VPS13, but not ERMES, is present, the Gem1 orthologue Miro was linked to mitochondria dynamics but not to lipid transport. Here we show that Miro, including its peroxisome-enriched splice variant, recruits the lipid transport protein VPS13D, which in turn binds the ER in a VAP-dependent way and thus could provide a lipid conduit between the ER and mitochondria. These findings reveal a so far missing link between function(s) of Gem1/Miro in yeast and higher eukaryotes, where Miro is a Parkin substrate, with potential implications for Parkinson’s disease pathogenesis.

**Summary:** VPS13D mutations result in severe mitochondrial defects. Guillén-Samander et al, show that VPS13D binds VAP in the ER, and interacts with Miro on mitochondria and peroxisomes, so that it can provide a bridge for lipid transport between these organelles.

---

Homeostasis of membranous subcellular organelles relies on appropriate synthesis, metabolism and distribution of bilayer lipids. Most such lipids are synthesized in the ER and their delivery to other destinations relies both on vesicular traffic and on protein-mediated lipid transfer. Much of the latter route of delivery occurs at membrane contact sites where many lipid transport proteins also function as tethers between the two membranes (Prinz, 2014; Saheki and De Camilli, 2016; Wong et al., 2019). For mitochondria, which are excluded from the membrane traffic flow of the secretory pathway, protein-mediated lipid transfer represents the only mechanism for their supply of lipids from the ER (Scharwey et al., 2013). In yeast, ERMES, a heterotetrametric protein complex that contains lipid transport modules and tethers the ER to the outer mitochondrial membrane (OMM) was shown to account for some of this transport(Kornmann et al., 2009). Another yeast protein, Vps13, which also localizes at membrane contact sites, albeit not ER-mitochondria contacts in this organism, was shown to have partially overlapping function with ERMES, possibly by participating in an alternative route for the delivery of lipids from the ER to mitochondria via the vacuole (Lang et al., 2015; Park et al., 2016; John Peter et al., 2017). While the ERMES complex is not conserved in metazoans, the single yeast Vps13 has 4 different homologues in mammals (including in humans)(Velayos-Baeza et al., 2004). Additionally, an accessory subunit of ERMES, the outer mitochondrial membrane GTPase Gem1, is also conserved in higher eukaryotes (Frederick et al., 2004; Kornmann et al., 2011). The two mammalian Gem1 orthologues, however, Miro1 and Miro2, have been associated primarily with mitochondrial dynamics and not with lipid transport (Fransson et al., 2003; Glater et al., 2006; Saotome et al., 2008; Macaskill et al., 2009; Wang and Schwarz, 2009; Nguyen et al., 2014). Notably, a localization of Miro at ER-mitochondria contacts sites has also been reported (Kornmann et al., 2009; Modi et al., 2019). Additionally, splice variants of Miro1 (Miro1v2 and Miro1v4) were shown to be enriched at peroxisomes and to be implicated in the dynamics of these organelles as well (Okumoto et al., 2018; Castro et al., 2018; Covill-Cooke et al., 2020).

Recent findings have shown that VPS13 is indeed a lipid transport protein and shed light on the molecular properties of VPS13 family proteins and on some of their sites of action in animal cells(Kumar et al., 2018; Li et al., 2020). The N-terminal portion of VPS13 folds as an elongated tube with a hydrophobic groove that runs along its entire length, thus suggesting that VPS13 acts as a bridge allowing lipid flow from one bilayer to another at sites of bilayer apposition (Li et al., 2020; Lees and Reinisch, 2020; Ugur et al., 2020). Such structure may account for net transfer of lipids and thus for a role of VPS13 and of its distant relative ATG2 (Kumar et al., 2018), an autophagy protein with a very similar fold(Valverde et al., 2019; Osawa et al., 2019; Maeda et al., 2019), in membrane expansion. Studies of mammalian VPS13A and VPS13C have demonstrated that they not only are localized at membrane contact sites, but also tether adjacent membranes(Kumar et al., 2018). Moreover, studies of the mammalian VPS13 protein family have shown paralogue-specific localizations (Seifert et al., 2011; Kumar et al., 2018; Yeshaw et al., 2019; Park and Neiman, 2020). VPS13A populates ER-mitochondria contacts, where it may have taken over some of the functions of ERMES, VPS13C is localized at ER-late endosomes/lysosome contacts(Kumar et al., 2018) and VPS13B resides predominantly in the Golgi complex region (Seifert et al., 2011). The localization and potential tethering properties of VPS13D remain unclear. Distinct functions of the four mammalian VPS13 proteins are further demonstrated by the different neurological diseases associated with their loss-of-function mutations, namely chorea acanthocytosis (VPS13A)(Rampoldi et al., 2001; Ueno et al., 2001), Cohen syndrome (VPS13B)(Kolehmainen et al., 2003), Parkinson’s disease (VPS13C)(Lesage et al., 2016; Schormair et al., 2018) and ataxia (VPS13D)(Gauthier et al., 2018; Seong et al., 2018).

VPS13D is the only mammalian VPS13 protein proven to be essential for cell and organismal survival (Wang et al., 2015; Blomen et al., 2015; Seong et al., 2018; Anding et al., 2018). While complete absence of each of the other three VPS13 proteins is compatible with life in humans, albeit with developmental (VPS13B) or neurodegenerative (VPS13A and VPS13C) defects, in patients with compound heterozygous VPS13D mutations leading to severe ataxias at least one of the alleles carries a missense mutation, thus probably encoding an at least partially functioning protein(Dziurdzik et al., 2020; Ugur et al., 2020). Complete absence of the VPS13D orthologue in mice leads to embryonic lethality, and in flies leads to death at the larval stage(Seong et al., 2018; Anding et al., 2018). Given its physiological importance, investigation of this protein will likely reveal some fundamental aspect of the biology of the VPS13 protein family. Studies in both human cells and flies have suggested a role for VPS13D in mitochondrial function. Enlarged spherical mitochondria have been reported in midgut cells and neurons of VPS13D knock-down flies, in VPS13D-KO HeLa cells and in patient-derived fibroblasts(Seong et al., 2018; Anding et al., 2018; Insolera et al., 2020 *Preprint*). However, so far, VPS13D has not been visualized in association with mitochondria, and was instead reported, based on immunofluorescence of its orthologue in Drosophila, to be localized on lysosomes(Anding et al., 2018).

Here we show that VPS13D binds to mitochondria and peroxisomes via an interaction with the GTPase Miro. It also binds the ER via its interaction with the ER protein VAP through a phospho-FFAT motif. These findings suggest that, like its yeast orthologue Gem1, Miro participates in the control of lipid transport between adjacent membranes, thus revealing an evolutionary link between the functions of Miro and Gem1 that had remained elusive until now. Collectively, our results provide new insight into the crosstalk between the ER and other organelles at contact sites and point to an explanation for the mitochondrial phenotypes caused by the lack of VPS13D.

## Results

### Localization of VPS13D in the Golgi complex and on mitochondria

VPS13D, with 4388 residues in humans, is the largest protein of the human VPS13 family. Its predicted domain architecture is overall very similar to that of other VPS13 proteins (Velayos-Baeza et al., 2004; Kumar et al., 2018). It comprises a long N-terminal lipid transfer domain that starts with the conserved chorein domain and is predicted to be slightly longer than in other VPS13 paralogues, a β-propeller region (so called VAB(Bean et al., 2018) or WD40-like region(Kumar et al., 2018)), a DH-like domain and a C-terminal PH domain. Differently from other human VPS13 proteins, VPS13D also contains a ubiquitin-binding domain (UBA) (Fig. 1A, left)(Anding et al., 2018; Ugur et al., 2020). To investigate VPS13D localization in cells, overcoming the absence of antibodies that yield a detectable immunofluorescence signal, and the limited expression expected upon transfection of such a large protein, a synthetic codon optimized cDNA sequence encoding VPS13D was generated, which also included a fluorescent tag (Halo or EGFP) after residue 1576 (hereafter referred to as VPS13D^tag). This position, which is localized in a predicted disordered loop, corresponds to the position where tag insertion did not affect function in other VPS13 proteins (Park et al., 2016; Kumar et al., 2018)(Fig. 1A, left). Expression of VPS13D^EGFP in COS7 cells resulted in a diffuse cytosolic fluorescence with some variable accumulation in the Golgi complex region and a weak enrichment, also variable in intensity from cell to cell, but detectable in about half of the cells, around mitochondria and in sparse additional spots (Fig. 1B). In cells where the localization on mitochondria was detectable, VPS13D was not restricted to a small subset of mitochondria, speaking against a selective binding of VPS13D via its UBA domain to ubiquitinated mitochondria targeted to degradation.

**Figure 1.**
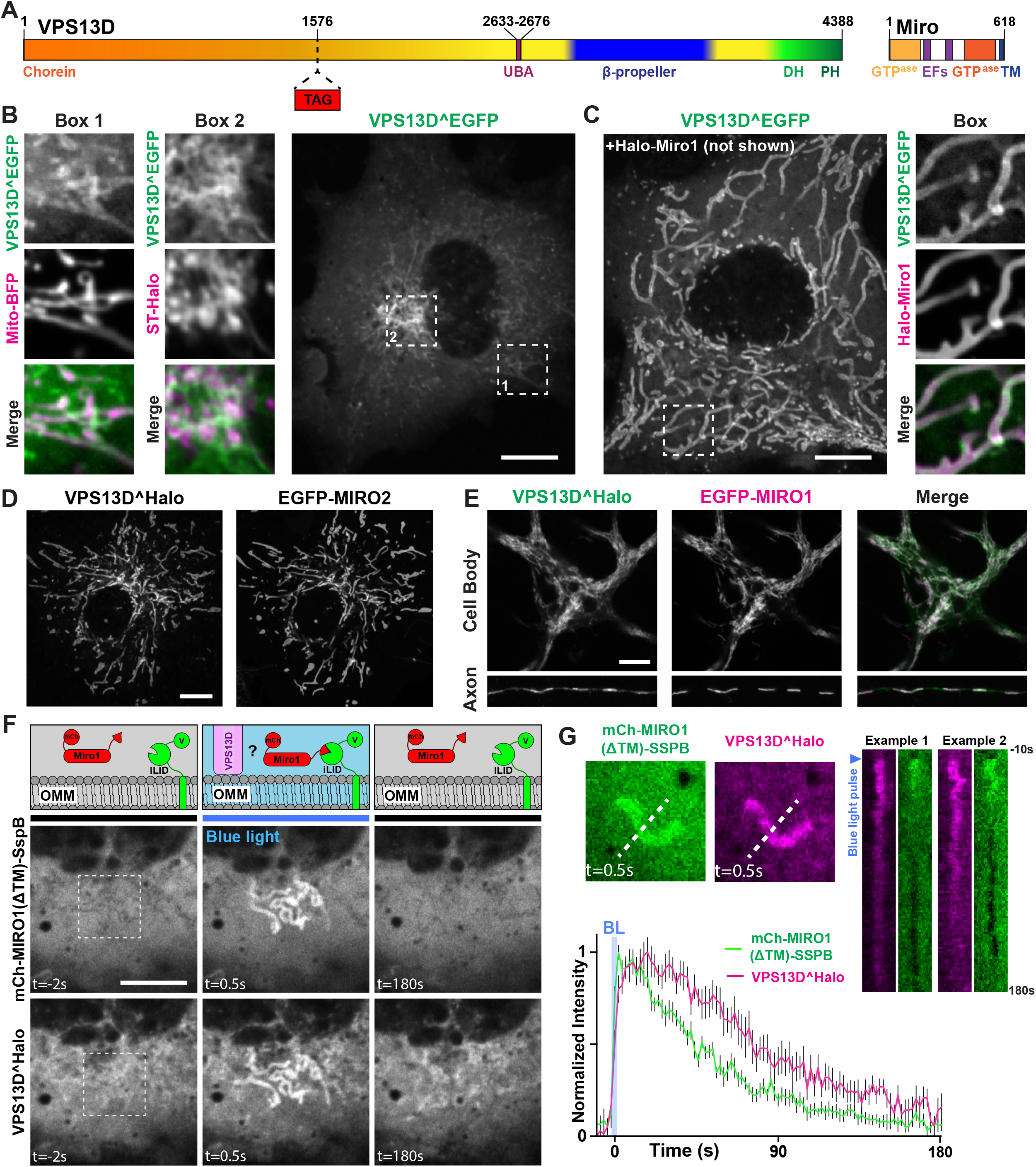
VPS13D recruitment by Miro GTPases. (A) Domain organization of human VPS13D and Miro1. (B) Confocal images of COS7 cell expressing VPS13D^EGFP showing that this protein decorates mitochondria, as shown by colocalization with mito-BFP (Box 1), in addition to being enriched in the Golgi area, visualized by the trans-Golgi marker Sialyltransferase(ST)-Halo (Box 2). (C) COS7 cell coexpressing VPS13D^EGFP and Halo-Miro1, showing dramatic increase in the localization of VPS13D^EGFP to mitochondria produced by Miro overexpression. The region within the white rectangle is shown at higher magnification next to the main field. (D) Recruitment of VPS13D to mitochondria can also be achieved by co-expression with Miro2. (E) VPS13D^Halo and EGFP-Miro1 colocalize in the cell body and processes of a mouse hippocampal neuron. (F) Optogenetic recruitment of the cytosolic domain of Miro1 to the outer mitochondrial membrane (OMM). Top panels: schematic representation of the experiment: Venus-iLID-Mito in the OMM recruits mCh-Miro1(ΔTM)-SspB upon blue-light excitation. The recruitment of Miro1, in turn, triggers the recruitment of VPS13D. Mid and bottom panels: confocal images of a COS7 cell showing blue-light-dependent recruitment and shedding of mCh-Miro1(ΔTM)-SspB and correspondingly of VPS13D^Halo. The illumination was started at time 0sec on a 5μm^2^ area shown in the leftmost panel. See also Movie S1. Scale bars=10μm. (G) Time-course of the recruitment and shedding of VPS13D^Halo and mCh-Miro1(ΔTM)-SSPB to the OMM. Top left panel: snapshots of an isolated mitochondrion at the peak of recruitment. Top right panel: example kymographs showing the increase and decrease in fluorescence along the stippled line shown in the left panels. Note that while the mCh-Miro1(ΔTM)-SSPB signal on mitochondria is lower than in the surrounding cytosol before illumination and after recovery, this is not the case for VPS13D^Halo, as a pool of this protein in bound to endogenous Miro. The kymographs start 10s before illumination. Bottom panel: Graph showing the normalized fluorescence intensity (average values ±SEM) along the length of kymographs of 18 independently illuminated mitochondria in 14 different cells. The decay in intensity of VPS13D^Halo and mCh-Miro1(ΔTM)-SspB on mitochondria was fitted to an exponential equation: For VPS13D^Halo, time constant τ=86.51s, 95%CI [77.64, 97.56], for mCh-Miro1(ΔTM)-SspB time constant τ=54.11s, 95%CI [51.07, 57.54], adj. R-square of fits 0.97 and 0.98, respectively.

### Overexpression of Miro very strongly enhances the recruitment of VPS13D to mitochondria

Among the outer mitochondrial membrane (OMM) proteins that could function as regulated binding partners for VPS13D at the mitochondrial surface, we considered Miro1 and Miro2 (henceforth referred to as Miro) as potential candidates. Miro, which contains two GTPase domains and two EF-hand Ca^2+^-binding domains exposed to the cytosol (Fig. 1A, right), has been primarily implicated in the regulation of mitochondrial transport in mammalian cells (Fransson et al., 2003; Glater et al., 2006; Saotome et al., 2008; Macaskill et al., 2009; Wang and Schwarz, 2009; Nguyen et al., 2014). The Miro orthologue in yeast, Gem1, is an accessory subunit of the ERMES complex, whose function in lipid transport between ER and mitochondria is partially redundant with that of yeast Vps13 (Lang et al., 2015; Park et al., 2016). Thus, a potential connection of Miro to a mammalian VPS13 family member seemed plausible. When either Miro1 or Miro2 were co-expressed with VPS13D in COS7 cells, a major impact on the localization of VPS13D was observed. This protein was now strikingly concentrated on mitochondria in more than 85% of the cells, where it colocalized with Miro along the entire mitochondrial surface (Fig. 1C and D and Fig. S1; see also Fig. 2B). Minimal residual fluorescence was observed elsewhere in the cell as if Miro had sequestered the entire VPS13D pool on mitochondria. Colocalization of co-transfected VPS13D^Halo and EGFP-Miro1 was also observed in mouse hippocampal neurons, where VPS13D was recruited to mitochondria both in the cell soma and in the neurites (Fig. 1E).

**Figure 2.**
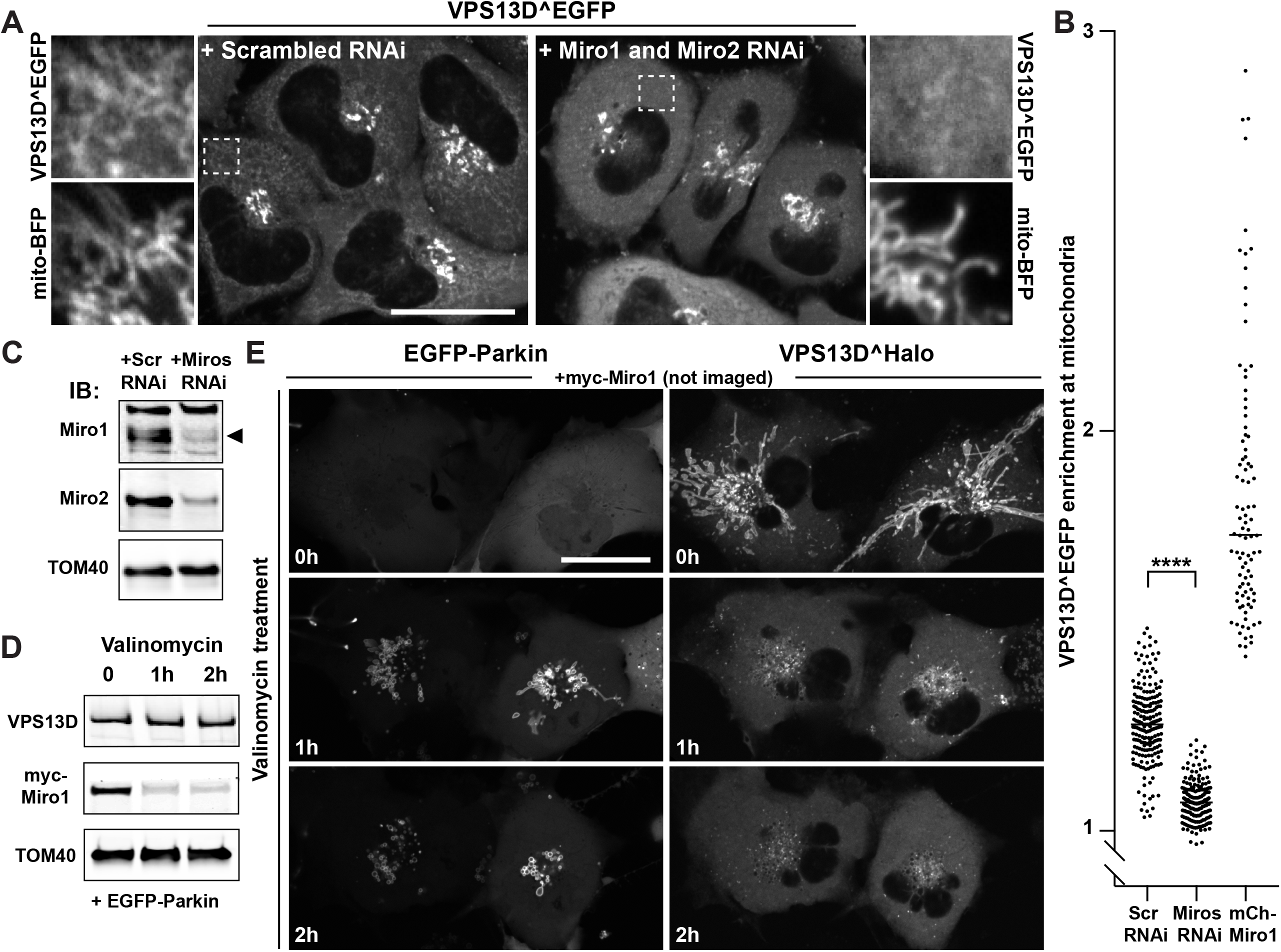
Miro is required for VPS13D recruitment to mitochondria. (A) Confocal images of HeLa cells showing the moderate recruitment of VPS13D^EGFP to mitochondria in control cells (left panel) and the loss of such signal upon knock-down of the two Miro genes via RNAi (right panel). Areas enclosed by stippled white rectangles are shown at higher magnification next to the main field, which also show the localization of the mitochondrial marker mito-BFP. Scale bars= 30μm. (B) Quantification of VPS13D^EGFP enrichment at mitochondria in control conditions and upon Miro knock-down or overexpression. The signal from mito-BFP was first used to generate a mitochondrial mask and a mask profiling a thin (1-pixel wide) cytosolic area surrounding mitochondria; the intensity from EGFP was then measured within each of these masks, and for each cell analyzed the ratio between these two measurements was plotted on the graph. \*\*\*\**P*<0.0001 (Welch’s corrected ANOVA, with Bonferroni post-hoc test) (C) Western blot of mitochondrial fractions from HeLa cells showing the decrease of Miro1 and Miro2 levels upon RNAi treatment. (D) Western blot showing the decrease of myc-Miro1 level upon 10μM Valinomycin treatment of COS7 cells co-transfected with EGFP-Parkin. (E) Confocal time lapse of COS7 cells co-expressing EGFP-Parkin, VPS13D^Halo and myc-Miro1. VPS13D, initially recruited to mitochondria by Miro, is shed from mitochondria upon treatment with Valinomycin, which induces the recruitment of EGFP-Parkin to initiate mitophagy. Scale bars= 30μm.

### Miro-dependent acute recruitment of VPS13D to mitochondria

To confirm a role for Miro in VPS13D recruitment to mitochondria, the iLID optical dimerization system(Guntas et al., 2015) was used to induce acutely, upon blue light illumination, the association of the cytosolic portion of Miro to the mitochondrial surface. We co-transfected VPS13D^Halo with a mitochondrial anchored bait (iLID fused to a mitochondrial targeting sequence) and the cytosolic portion of Miro1 fused to the cognate pray for iLID (the SspB peptide). Blue light illumination induced immediate massive recruitment of the Miro1 construct to mitochondria which was rapidly followed by a corresponding massive recruitment of VPS13D (Fig. 1F and 1G), proving the role of Miro in VPS13D recruitment. Conversely, upon light interruption, both proteins were shed from mitochondria (Fig. 1G and Movie S1). The short latency of the shedding of VPS13D relative to shedding of Miro during the post-illumination phase (Fig. 1G), may reflect the occurrence of stabilizing VPS13D interactions at the mitochondrial membrane, including its interaction with endogenous Miro, once its recruitment has occurred.

### Loss of Miro results in a defect in the targeting of VPS13D to mitochondria

We next examined whether Miro is required for the recruitment to mitochondria of exogenous VPS13D. HeLa cells were transfected with VPS13D^EGFP and short interfering RNAs (siRNAs) targeting the two Miro genes or control scrambled siRNAs. As shown by Fig. 2A, and quantified in Fig. 2B, in cells expressing Miro siRNAs, where the decrease of both Miro1 and Miro2 was validated by western blotting (Fig. 2C), the VPS13D^EGFP signal on mitochondria was significantly reduced, while the robust signal in the Golgi complex, where Miro is not localized, was not affected. We also attempted to analyze the localization of VPS13D^EGFP in cells where one Miro paralogue was knocked-out by CRISPR-Cas9 and the other one was knocked down, but the poor viability of these cells prevented a reliable analysis.

Miro is a key target of the PINK1/Parkin-dependent degradation pathway of mitochondrial proteins leading to mitophagy. Triggering of this pathway with valinomycin, which disrupts mitochondrial membrane potential and promotes the recruitment of Parkin to mitochondria (Narendra et al., 2010; Yamano et al., 2018), results in the rapid degradation of Miro proteins (Wang et al., 2011; Birsa et al., 2014; Shlevkov et al., 2016). Accordingly, 1h of valinomycin treatment of either HeLa or COS-7 cells expressing myc-Miro1 resulted in a robust decrease in the levels of this protein (Fig. 2D and Fig. S2A). While levels of VPS13D, either endogenous (Fig. 2D) or exogenous (VPS13D^mScarlet, Fig. S2A) did not change, Miro degradation coincided with the shedding of VPS13D^Halo from mitochondria and with an increase in its cytosolic pool (Fig. 2E and S2B). These results establish the requirement of Miro in the recruitment of VPS13D to mitochondria.

### A Miro1 splice variant targeted to peroxisomes recruits VPS13D to peroxisomes

A splice variant of Miro including exons #19 and #20 in its C-terminal region (transcript variant 4, referred to as Miro1v4) was shown to localize predominantly at the membrane of peroxisomes, and only weakly on the OMM, via a mechanism that requires its interaction with Pex19(Okumoto et al., 2018; Covill-Cooke et al., 2020)(Fig. 3A), a cytosolic chaperone that functions as a receptor for a subset of peroxisomal membrane proteins(Jones et al., 2004). If Miro is sufficient to recruit VPS13D to an organelle, one would expect that Miro1v4 may recruit VPS13D to peroxisomes. In fact, upon co-expression with Miro1v4 in COS7 cells, the majority of VPS13D^Halo was recruited to peroxisomes where Miro1v4 was predominantly localized, and only a faint VPS13D^Halo signal was observed on mitochondria (Fig. 3B).

**Figure 3.**
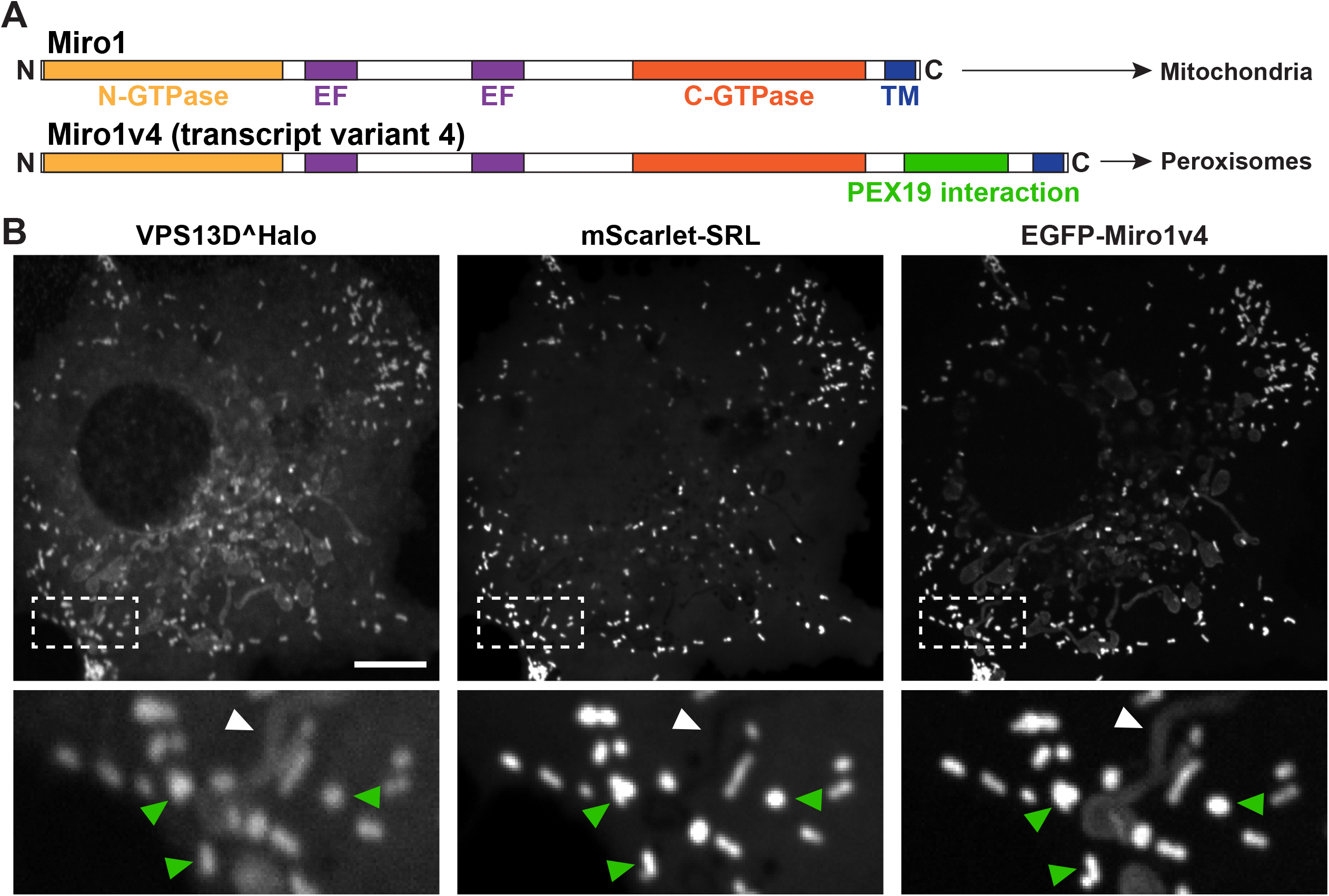
A transcript variant of Miro1 preferentially targeted to peroxisomes recruits VPS13D to peroxisomes. (A) Domain organization of two splice variants of Miro1(Okumoto et al., 2018). In the alternative splice variant 4 (Miro1v4) two extra exons are included that encode 73 additional aminoacids in the C-terminal region of Miro1. These aminoacids promote the interaction of the protein with the chaperone Pex19, which leads to the predominant insertion of this variant into the peroxisomal membrane, although a pool of this variant still localizes to mitochondria. (B) COS7 cells recruitment of VPS13D^Halo to peroxisomes (visualized by the peroxisomal luminal marker mScarlet-SRL) upon co-expression with transcript variant Miro1v4. The insets show colocalization with the peroxisomal marker (green arrowheads) and a weaker signal for both VPS13D and Miro1v4 on mitochondria (white arrowhead). Scale bars= 10μm.

### Integrity of the GTPase domains and EF hand regions of Miro are required for the recruitment of VPS13D

Binding of Miro1 to some of its known other interactors and of Gem1 to ERMES in yeast requires integrity of its GTPase and EF hand domains(Kornmann et al., 2011; Kanfer et al., 2015; Oeding et al., 2018). When inactivating mutations based on previous studies were introduced in the GTP binding site of N-terminal GTPase domain (T18N) or in the Ca^2+^-binding sites of both EF hands (E208K/E328K) of Miro1(Fransson et al., 2003; Frederick et al., 2004) (Fig. 4A), the corresponding constructs no longer promoted recruitment of VPS13D to mitochondria (Fig. 4B). In contrast, a mutation in the C-terminal GTPase (S432N) of Miro1(Fransson et al., 2003; Frederick et al., 2004) did not affect the recruitment of VPS13D (Fig. 4B). To assess a potential regulatory role of cytosolic Ca^2+^, we treated cells with the ER Ca^2+^ pump inhibitor thapsigargin to raise its concentration, but no further increase of VPS13D recruitment to mitochondria in cells overexpression Miro1 was observed (Fig. S3A and S3B). Likewise, incubation of cells in the absence of extracellular Ca^2+^ and in the presence of EGTA (to chelate residual extracellular Ca^2+^) and BAPTA-AM, to chelate intracellular Ca^2+^, had no obvious effect in VPS13D recruitment to mitochondria (Fig. S3C). In light of this insensitivity to cytosolic Ca^2+^, the possibility that some of the disrupting effects of these mutations reflect regulatory mechanisms or protein misfolding, as previously considered(Kanfer et al., 2015), remains to be determined.

**Figure 4.**
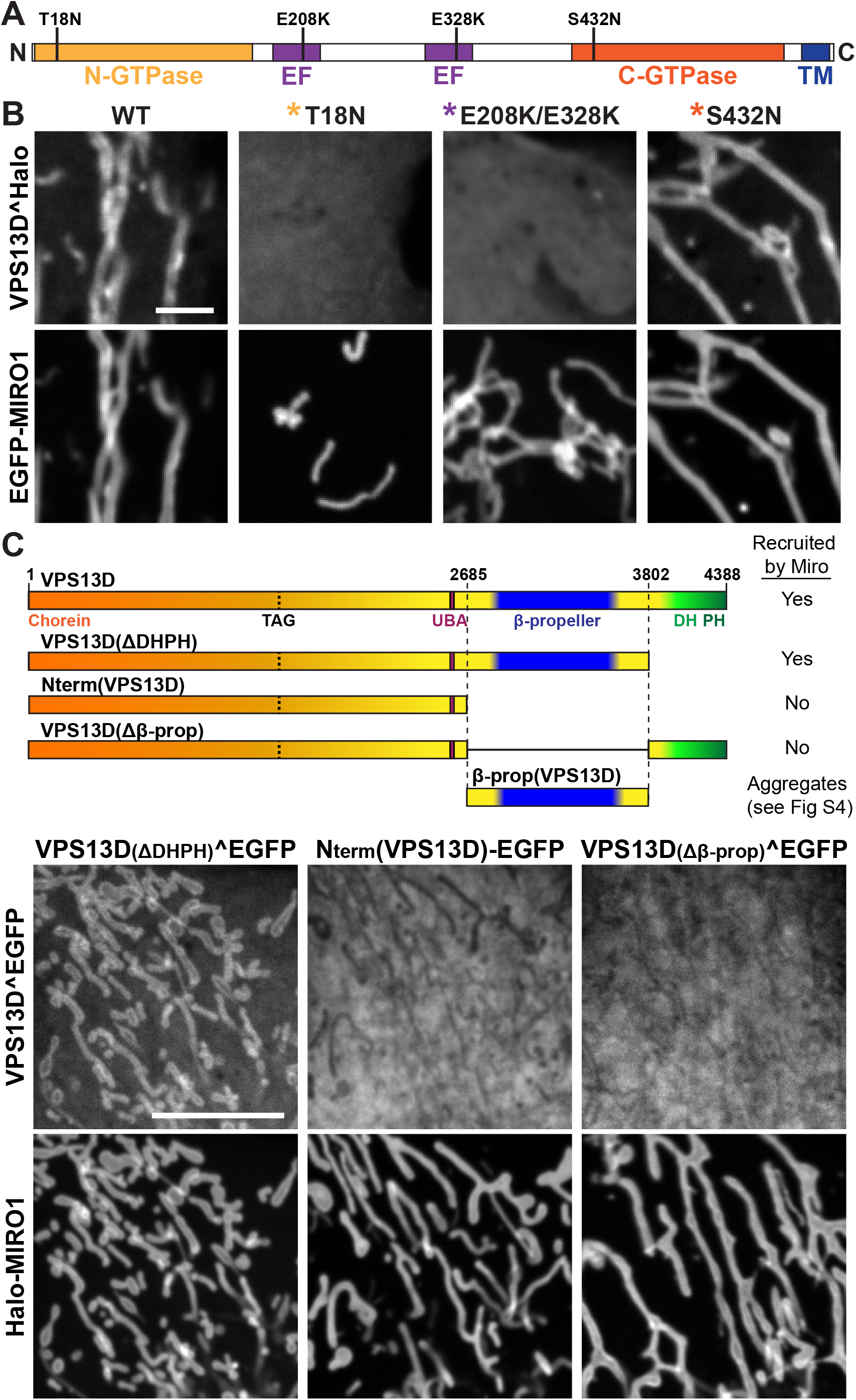
Recruitment of VPS13D by Miro GTPases requires their intact N-GTPase domain and EF-hands domains, and the β-propeller region of VPS13D. (A) Domain organization of human Miro1 and mutations used for the experiments of panel B. (B) Confocal images of COS7 cells showing that the recruitment of VPS13D^Halo by EGFP-Miro1 is impaired by mutations in the GTP binding site of its N-GTPase domain and in the Ca^2+^ binding sites of both its EF-hands, but not by a mutation in the GTP binding site of its C-GTPase domain. Scale bars=3μm. (C) Top: Cartoons showing VPS13D constructs used for the experiments shown below. Bottom: Confocal images showing that removal of the C-terminal half of the protein or of the β-propeller region selectively, but not of the DHPH domain, abolishes recruitment of VPS13D^EGFP by Halo-Miro1. All of several constructs encoding only the β-propeller region formed small aggregates, possibly due to misfolding. See also Fig. S4. Scale bars=10μm

### The VAB/β-propeller domain of VPS13D is required for its recruitment by Miro

To determine the region of VPS13D required for Miro-dependent recruitment, we co-expressed in COS7 cells Miro with several VPS13D fragments. Removal of the C-terminal region of VPS13D, including both the VAB/β-propeller domain and DH-PH domains, abolished its Miro-dependent recruitment to mitochondria (Fig. 4C). In contrast, VPS13D constructs lacking the DH-PH domain only were still recruited by Miro, whereas constructs containing the DH-PH domain but not the VAB//β-propeller domain were not (Fig. 4C), suggesting that the VAB/β-propeller domain was necessary for this recruitment. Efforts to determine whether the VAB/β-propeller domain was sufficient for the recruitment were inconclusive, as each of all of several constructs of this domain that we generated produced heterogenous subcellular localizations ranging from cytosolic distribution, to small aggregates, to a colocalization with mitochondria (with roughly equal proportions of the three localization patterns). However, in the latter case the morphology and intracellular distribution of mitochondria was disrupted, often resulting in their massive clustering, as shown by the EGFP-Miro1 signal. We interpret these results as suggesting that VAB/β-propeller alone may not fold correctly (Fig. S4). While all of our data are compatible with a direct interaction between VPS13D and Miro, we cannot rule out an indirect interaction.

VPS13A, a paralogue of VPS13D, also binds mitochondria, but does so via its C-terminal DH-PH domain whose interactor(s) at the mitochondrial surface remain(s) unknown(Kumar et al., 2018). Accordingly, co-transfection of VPS13A^Halo with EGFP-Miro1 did not result in an increased recruitment of VPS13A^Halo to mitochondria (Fig. S1C-E). Moreover, when VPS13A^Halo and VPS13D^EGFP were cotransfected in COS7 cells (Fig. S1E) also overexpressing Miro1, they were both concentrated at mitochondria but with a different pattern: VPS13A had a punctate distribution previously shown to reflect its restricted localization at ER-mitochondria contact sites(Kumar et al., 2018), while VPS13D decorated the entire mitochondrial surface. This indicates a different mechanism of mitochondrial binding of VPS13A and VPS13D.

### VPS13D binds the ER protein VAP via an unconventional FFAT motif

Studies of yeast Vps13 and of mammalian VPS13A and VPS13C have shown that a shared property of these proteins is to bridge organelles to provide a conduit for the flow of lipids between adjacent bilayers. Both VPS13A and VPS13C tether other organelles to the ER via an interaction of their FFAT motifs with the MSP domain of the ER proteins VAP-A and VAP-B (Kumar et al., 2018).While the homogenous localization of VPS13D on the mitochondrial surface does not support a selective localization of this protein at contact sites with the ER, we considered that an interaction of VPS13D with the ER, and possibly with VAP, may be of low affinity and/or regulated. To explore this possibility, we examined the localization of VPS13D^EGFP in COS7 cells when co-expressed with Halo-VAP-B but without Miro. While no obvious fluorescence signal on the ER was observed when VPS13D^EGFP was expressed alone (Fig. 1B), a weak ER localization of VPS13D^EGFP was observed in cells also expressing Halo-VAP-B (Fig. 5A). Most likely, endogenous VAP expression levels are not sufficient to bind a detectable amount of VPS13D and/or the diffuse cytosolic fluorescence of VPS13D^EGFP obscures such localization. Overexpression of VAP-B, but not of a VAP-B construct harboring two mutations (K87D/M89D) that impair FFAT motif binding(Dong et al., 2016), also induced the recruitment to the ER of the VPS13D N-terminal region alone, further validating VAP-dependent binding, and indicating that the ER binding site, as in the case of VPS13A and VPS13C, lies somewhere in this region (Fig. 5B and 5E).

**Figure 5.**
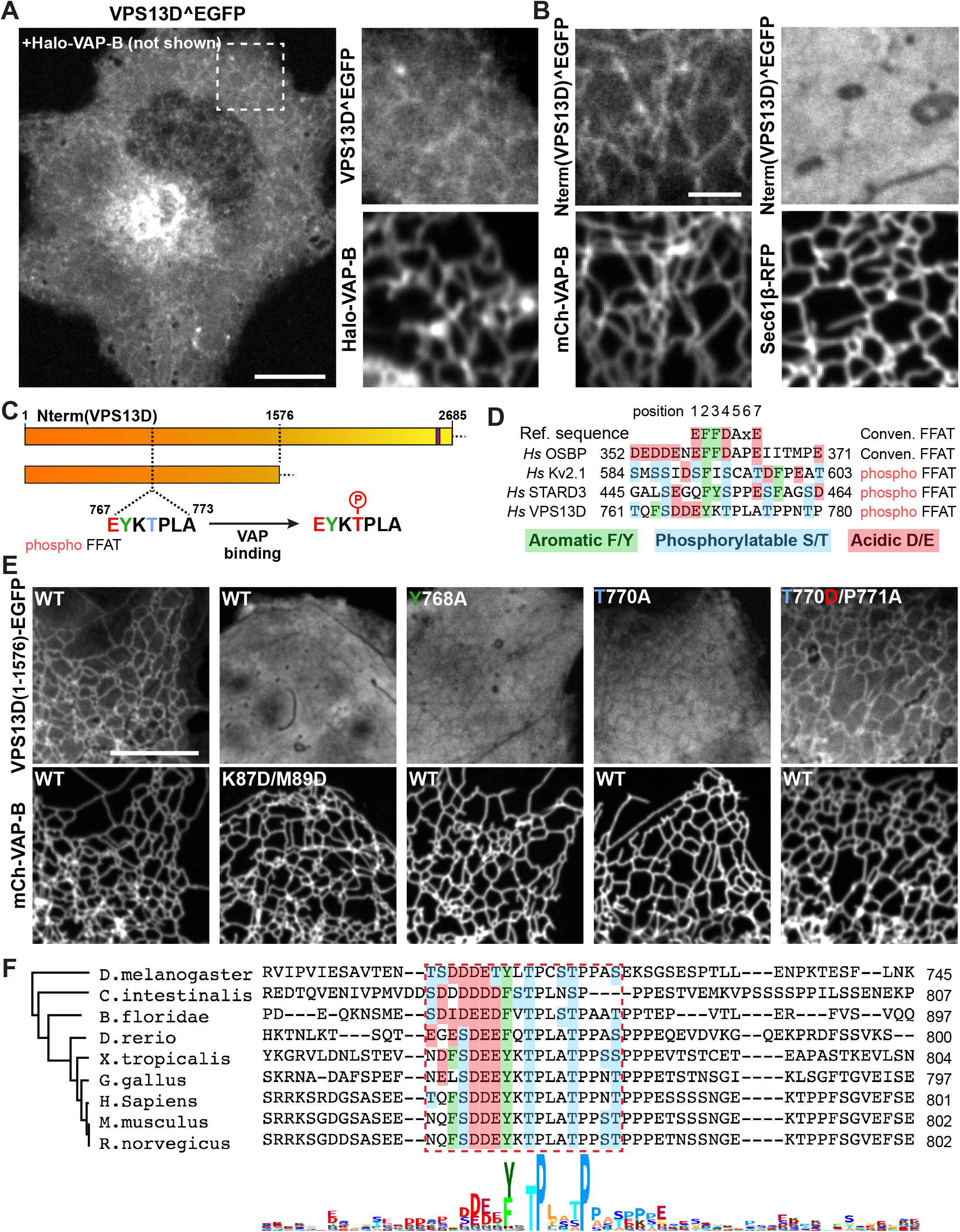
VPS13D binds VAP on the ER via a phospho-FFAT in its N-terminal region. (A) Left: confocal image of a COS7 cell co-expressing Halo-VAP-B and VPS13D^EGFP showing weak recruitment of VPS13D^EGFP to the ER. Scale bar=10μm. Right: high magnification view of Halo-VAP-B and VPS13D^EGFP fluorescence of the field enclosed by a rectangle. (B) VAP-dependent binding of the N-terminal portion of VPS13D to the ER. Left: confocal images of a COS7 cell co-expressing the N-terminal portion of VPS13D fused to EGFP and mCherry-VAP-B. Right: confocal images of a COS7 cell co-expressing the N-terminal portion of VPS13D fused to EGFP and Sec61β-RFP, but not VAP. Scale bar= 3μm (C) Cartoons showing VPS13D constructs used for the experiments shown in Fig 5E, indicating the position of the predicted phospho-FFAT motif. (D) Comparison of the conventional FFAT motif with phospho-FFAT motifs, including the one found in VPS13D(Di Mattia et al., 2020). The aromatic residue indicated in green, present in both conventional and phospho-FFAT motifs, is essential for binding to VAP. Acidic aminoacid residues in the region contribute to the binding in conventional FFAT motifs, but, based on a previous study (Di Mattia et al., 2020), they can replaced by phosphorylatable residues in phospho-FFAT motifs. (E) Evidence for a phospho-FFAT motif-dependent binding of the N-terminal region of VPS13D to VAP. First column: the N-terminal construct (1-1576) of VPS13D is recruited to the ER upon VAP overexpression. Second column: two mutations in the MSP domain of VAP that disrupt the FFAT motif binding pocket also disrupt the recruitment of the VPS13D construct. Third column: Mutation to alanine of the aromatic residue of the phospho-FFAT motif disrupts binding. Fourth and fifth column: no binding occurs when the threonine that corresponds to an aspartate in the conventional FFAT motif is replaced by a non-phosphorylatable alanine, but binding is restored when the threonine is replaced by aspartate, as long as the adjacent proline is also mutated(Di Mattia et al., 2020). Scale bar= 10μm (F) Alignment of the region of VPS13D orthologues from different species centered around the amino acid region required for Miro binding in human VPS13D. The alignment shows a high degree of conservation of the key residues of the phospho-FFAT motif among several chordates and also observed in flies. The phylogenetic tree was generated by maximum-likelihood.

Three conventional FFAT motifs are predicted with low score(Murphy and Levine, 2016; Slee and Levine, 2019) in the N-terminal region of VPS13D, but, surprisingly, the combined mutation of all three sites did not abolish VAP-dependent recruitment to the ER (Fig. S5). Recent studies of VAP binding proteins revealed the occurrence of unconventional VAP binding motifs that bind the FFAT binding pocket of VAP in spite of several differences, including the proposed dependence on phosphorylation rather than on acidic amino acids (hence referred to as phospho-FFAT motifs) (Johnson et al., 2018; Kirmiz et al., 2018; Di Mattia et al., 2020). As a phospho-FFAT motif was predicted in the N-terminal region of VPS13D (a.a. 767-773) (Di Mattia et al., 2020) (Fig. 5C and 5D), we tested its importance. Mutation to alanine of tyrosine (Y768A) at position 2 of the core motif, i.e. a position where there is an absolute requirement for an aromatic amino acid in all FFAT motifs, abolished the recruitment of the N-terminal region of VPS13D to the ER in VAP-B overexpressing cells (Fig. 5E, third column). Furthermore, mutation to alanine of the threonine (T770A) of the motif, which was suggested as a phosphorylatable site (Di Mattia et al., 2020), also disrupted the interaction with VAP (Fig. 5E, fourth column). Conversely, a phosphomimetic mutation of this threonine to aspartate, when combined with the mutation of the adjacent proline to alanine (T770D/P771A) rescued its ability to bind VAP (Fig. 5E, fifth column). The proline residue immediately downstream of the phosphosite is a shared characteristic among other phospho-FFAT motifs, and mutation of this proline to alanine was shown to be necessary to allow for the phospho-mimetic construct to bind VAP(Di Mattia et al., 2020). It was suggested by Di Mattia *et al*(Di Mattia et al., 2020) that the proline “would prevent the aspartate residue from properly mimicking a phosphorylated serine, as it possesses a shorter side chain”. Alignment of VPS13D protein sequences from different animal species revealed a high conservation of this phospho-FFAT motif, pointing to its physiological importance (Fig. 5F).

### Binding of VPS13D to VAP and to Miro allows it to function as a bridge between the ER and mitochondria

Prompted by the property of the N-terminal region of VPS13D to bind VAP, we examined whether VPS13D can bridge ER to mitochondria. When VPS13D^Halo was co-expressed with both BFP-VAP-B and EGFP-Miro1, the signal of VPS13D^Halo no longer decorated the entire mitochondrial surface homogeneously but was primarily concentrated at hot spots. Such hot spots precisely colocalized with sites of apposition between the mitochondria and the ER, consistent with a localization of VPS13D at contacts between the two organelles (Fig. 6A). This conclusion was supported by observing VPS13D dynamics during exposure of cells to strong hypotonic solutions (Fig. 6B and Movie S2). Under these conditions, dramatic shape changes and vesiculation of organelles occurs, but tethers between organelles persist(King et al., 2020). In cells expressing all three proteins, VPS13D^Halo fluorescence coalesced at sites where mitochondria- and ER-derived vesicles remained in contact with each other, as expected for a protein that can bridge the two organelles (Fig. 6B and Movie S2).

**Figure 6.**
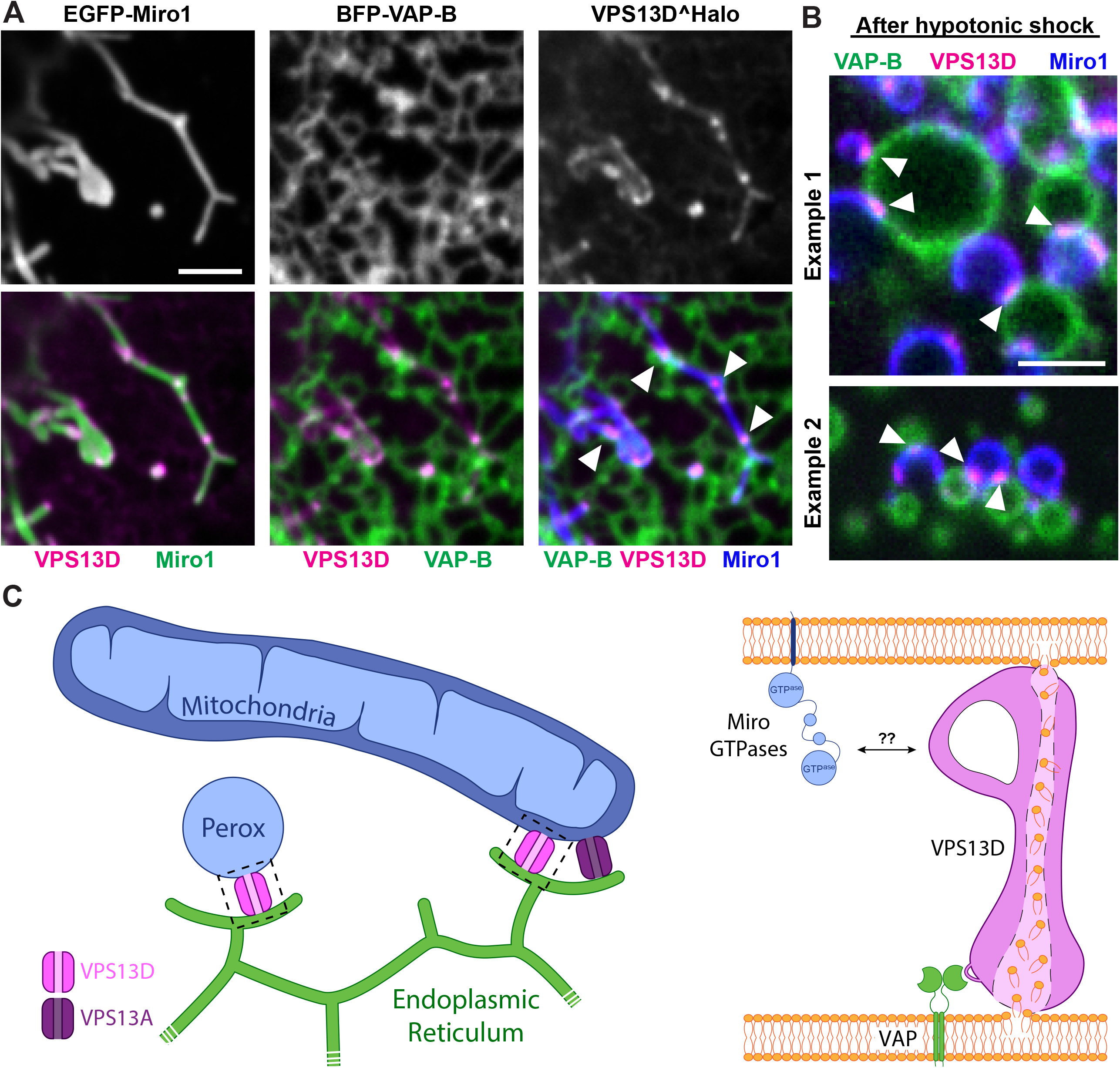
VPS13D can tether the ER to mitochondria in a VAP and Miro-dependent way. (A) COS7 cells co-expressing EGFP-Miro1, BFP-VAP-B and VPS13D^Halo. Top: single fluorescence images. Bottom: merge of the different fluorescence channels as indicated. White arrowheads in the triple merge shows that VPS13D^Halo (magenta) is enriched at hot spots where the ER (green) and mitochondria (blue) intersect. (B) Vesiculated ER and mitochondria induced by hypotonic treatment of COS7 cells co-expressing VPS13D^Halo, GFP-Miro1 and BFP-VAP-B remain tethered to each other, and VPS13D^Halo concentrates at these sites (white arrowheads). See also Movie S2. Scale bars=3μm. C: Schematic cartoon summarizing key findings of this study. Left: VPS13D can bridge the ER and either mitochondria or peroxisomes. Right: the interaction of VPS13D with the ER is mediated by VAP and its interaction with either mitochondria or peroxisomes is mediated by Miro. Based on the reported properties of VPS13 family proteins, it is proposed that VPS13D allows flux of lipids between the two tethered bilayers.

## Discussion

Our findings provide insight into the properties of VPS13D, a VPS13 family member previously shown to have an impact on mitochondrial biology based on loss-of-function studies, but whose site of action remained elusive. We show that VPS13D can bridge mitochondria to the ER and identify proteins on the two organelles responsible for this localization, the protein VAP in the ER membrane and the protein Miro in the outer mitochondrial membrane (Fig. 6C). Splice variants of Miro are targeted to peroxisomes(Okumoto et al., 2018; Covill-Cooke et al., 2020) and, accordingly, we have shown that VPS13D can also be recruited to peroxisomes. A membrane bridging property of VPS13D is consistent with the recently discovered core function of VPS13 proteins: mediating lipid fluxes between bilayers(Kumar et al., 2018; Li et al., 2020). Both mitochondria and peroxisomes, whose functions are closely interrelated, critically depend on non-vesicular lipid transport from the ER for their growth and function(Vance, 1990; Raychaudhuri and Prinz, 2008; Kornmann et al., 2009; Petrungaro and Kornmann, 2019).

An association of VPS13D with mitochondria is also supported by recent proximity-labelling based studies listing this protein among the hits retrieved with outer membrane protein baits (Hung et al., 2017; Liu et al., 2018; Antonicka et al., 2020). Interestingly, these studies also retrieved VPS13A, which we had shown previously to act as bridge between ER and mitochondria(Kumar et al., 2018), but did not retrieve VPS13B or VPS13C, for which there is no evidence for such a localization(Seifert et al., 2015; Kumar et al., 2018; Ugur et al., 2020). Based on these results, VPS13D may have a partially overlapping role with VPS13A in allowing lipid fluxes between the ER and mitochondria. However, VPS13D likely has a more fundamental function, as it is required for life(Wang et al., 2015; Blomen et al., 2015; Seong et al., 2018), while VPS13A is not. Clearly, the organelle bridging roles of VPS13A and VPS13D are differentially regulated, as the recruitment of VPS13A to mitochondria is not mediated by Miro (the mechanism of its binding to mitochondria remains unknown) and the FFAT motifs through which the two proteins bind VAP have different properties.

While VPS13A has a conventional FFAT motif(Murphy and Levine, 2016; Kumar et al., 2018), the VAP binding site of VPS13D fits the consensus of the recently defined phospho-FFAT motif(Di Mattia et al., 2020), so called because acidic amino acids of the motif can be replaced by phosphorylatable serines or threonines. In the phospho-FFAT motif of VPS13D, position 2 of the core sequence is occupied by a tyrosine (Y768), which we have shown to be essential for VAP binding as predicted, and position 4 is occupied by a threonine (T770) instead of an acidic amino acid as in conventional FFAT motifs. We have shown that replacement of this threonine with the non-phosphorylatable alanine abolishes binding, while replacement with the phosphomimetic aspartate rescues binding. Additional serines/threonines are present in the proximity of the core phosphoFFAT motif of VPS13D which may contribute to binding affinity via phosphorylation, as has been shown to happen in other phosphoFFAT motifs (Di Mattia et al., 2020).

The different regulation of the bridging functions of VPS13A and VPS13D is further revealed by the observation that, at least in the cells used in this study, detectable ER binding of VPS13D, but not of VPS13A, requires VAP overexpression, suggesting a lower affinity, likely regulated by phosphorylation. As a result of this difference, in cells that do not overexpress VAP, overexpressed VPS13D localizes along the entire mitochondrial surface, while overexpressed VPS13A accumulates at hot spots, which represent sites of ER-mitochondria contacts(Kumar et al., 2018)(Fig. S1).

The recruitment of VPS13D to peroxisomes via a splicing variant of Miro1 points to VPS13D as a critical player in delivering lipids to these organelles from the ER and is consistent with previous data indicating the importance of peroxisome-ER contacts for the expansion of their membranes in fungi(Raychaudhuri and Prinz, 2008; Akşit and van der Klei, 2018; Islinger et al., 2018) and more specifically of Vps13 in *H. polymorpha*(Akşit, 2018). Roles of VPS13D at other cellular sites should also be considered, in view of the enrichment of VPS13D^EGFP in the Golgi complex area, a localization that we have not further investigated in this study. Interestingly, another VPS13 paralogue, VPS13B, is selectively enriched in the Golgi complex area (Seifert et al., 2011; 2015), once again suggesting differences, but also partial overlap in the role of distinct VPS13 paralogues.

A partnership between VPS13D and Miro fits with the critical role of both proteins for life (López-Doménech et al., 2018; Seong et al., 2018). Moreover, in cellular models, loss-of-function perturbations of either Miro or VPS13D include similar alterations in mitochondrial size, distribution and degradation(Nguyen et al., 2014; Anding et al., 2018; Seong et al., 2018; López-Doménech et al., 2018; Insolera et al., 2020 *Preprint*). The accumulation of round mitochondria is a particularly striking feature of VPS13D and Miro deficient cells. Among the many potential scenarios to be considered, one is that the altered mitochondrial morphology in cells lacking VPS13D may result from reduced membrane lipid availability for membrane expansion with an increase of volume to surface ratio. Swelling of mitochondria, in turn, may impair the action of the fission machinery. Interestingly, it was reported that in mammalian cells expansion of the so-called “mitochondrial derived compartment”, a newly described membranous organelle derived from the outer mitochondrial membrane, requires Miro and contacts between mitochondria and the ER, consistent with a role of Miro in a process that may involve lipid flux from the ER (English et al., 2020; Schuler et al., 2020 *Preprint*).

Collectively, our study advances our understanding of pathways of non-vesicular lipid traffic within cells. It adds new evidence for the fundamental importance in cell physiology of proteins of the VPS13/ATG2 superfamily, i.e. proteins thought to act in the bulk delivery of lipids from one bilayer to another to mediate bilayer expansion(Ugur et al., 2020; Lees and Reinisch, 2020). It also reveals an evolutionary link, missing so far, between the function of the Gem1/Miro family in fungi, where Gem1 is an accessory subunit of ERMES(Kornmann et al., 2011), a major lipid transfer complex, and in mammals, where Miro had been primarily implicated in the control of mitochondrial dynamics and/or transport via molecular motors(Fransson et al., 2003; Saotome et al., 2008; Wang and Schwarz, 2009; Nguyen et al., 2014; López-Doménech et al., 2018). While in yeast the single Vps13 can compensate for the lack of the ERMES complex, no link between Gem1 and Vps13 has been observed so far in this organism. But ERMES is not present in metazoa, possibly explaining why the role of Gem1 as a partner of ERMES in lipid transport has evolved to a role of its orthologue Miro as a partner of VPS13D in mammalian cells and possibly other high eukaryotes. It will be of interest in the future to determine how the distinct functions of Miro in the control of mitochondria mobility and lipid transport are coordinated.

Finally, as Miro is a substrate for Parkin(Wang et al., 2011; Birsa et al., 2014; Shlevkov et al., 2016), and is itself a target of Parkinson’s Disease-associated mutations(Berenguer-Escuder et al., 2019), our results may also be relevant to pathogenetic mechanisms of Parkinson’s diseases.

## Supplementary Figure Legends

**Figure S1.**
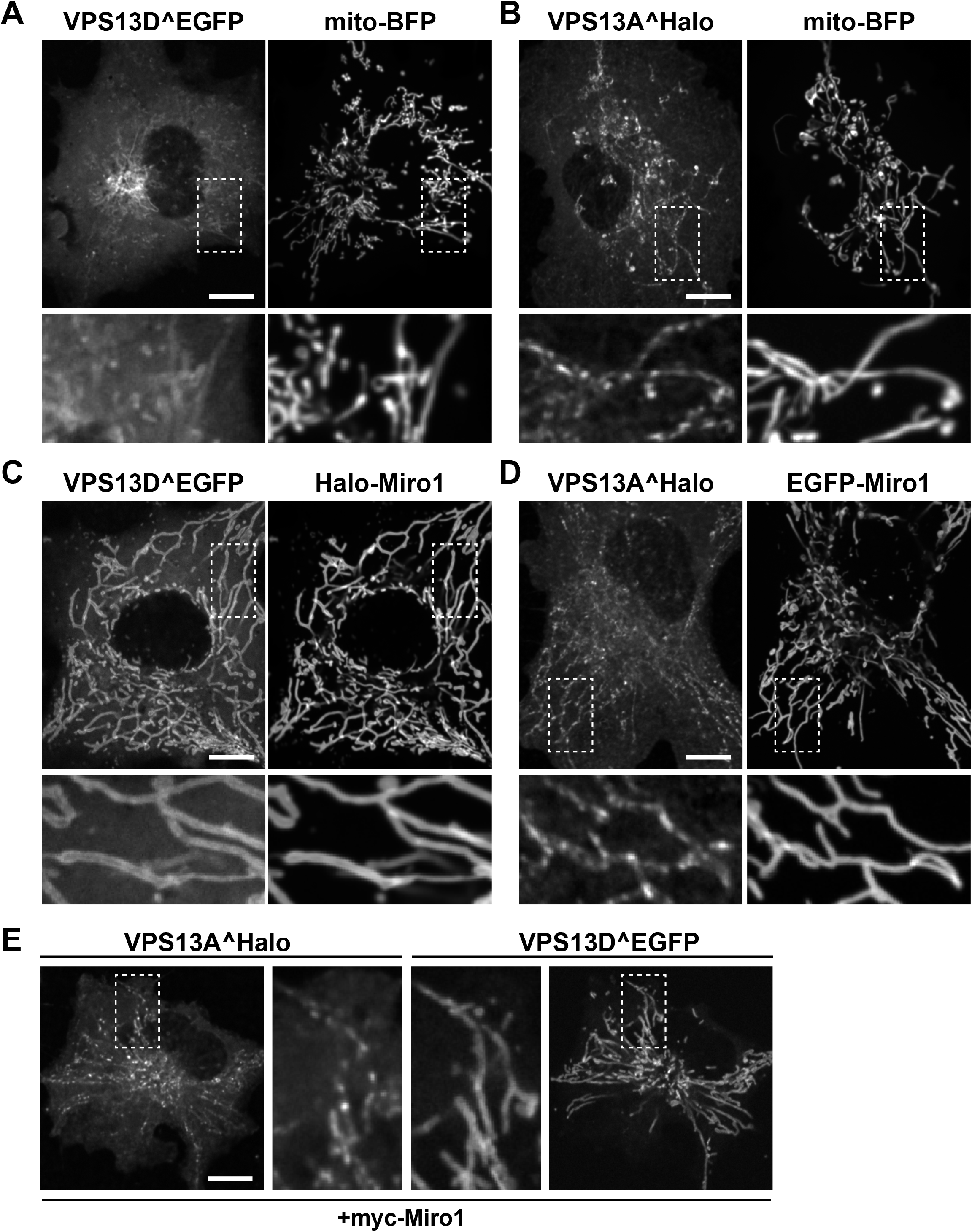
Both VPS13D and VPS13A interact with mitochondria, but only the interaction of VPS13D is mediated by Miro. (A and B) Confocal images of COS7 cells co-expressing either VPS13D^EGFP (A) or VPS13A^Halo (B) and the mitochondrial marker mito-BFP, but not Miro. The fluorescence of both VPS13 paralogues decorate mitochondria, with the fluorescence of VPS13A showing the typical discontinuous pattern which was shown(Kumar et al., 2018) to reflect its selective concentration at mitochondria-ER contact sites even without VAP overexpression. (C and D). Upon co-expression with Miro1, the localization of VPS13D^EGFP (C) but not VPS13A^Halo (D) at mitochondria is drastically enhanced. Note that VPS13D^EGFP colocalizes with Miro1 along the entire mitochondrial surface, while the punctate localization of VPS13A^Halo along mitochondria is unaffected by the overexpression of Miro1. (E) COS7 cell coexpressing VPS13A^Halo and VPS13D^EGFP as well as myc-Miro1(not shown), showing the different localization of the two proteins on mitochondria: VPS13D^EGFP is enriched throughout the entire mitochondrial surface, while VPS13A^Halo localizes only to hotspots. Higher magnifications of the areas enclosed by stippled white rectangles are shown for all fields. Scale bars= 10μm.

**Figure S2.**
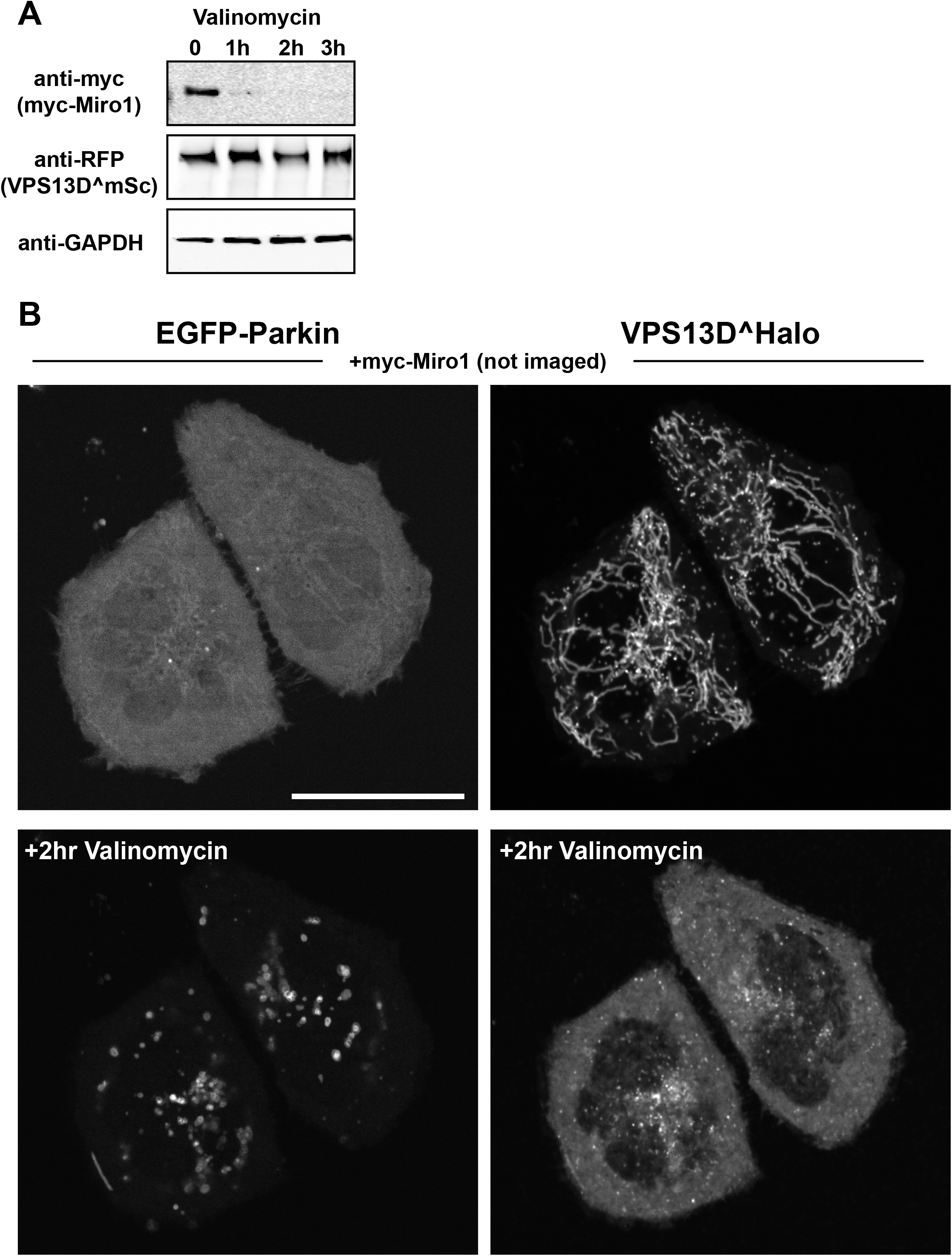
Parkin-mediated Miro degradation results in dissociation of VPS13D from mitochondria. HeLa cells were co-transfected with myc-Miro1, VPS13D^Scarlet and EGFP-Parkin. (A) Western blot showing the decrease in levels of myc-Miro1, but not of VPS13D^Scarlet, upon Valinomycin treatment. Scarlet, a modified RFP, was visualized by anti-RFP antibodies. (B) Confocal time lapse of HeLa cells co-expressing EGFP-Parkin, VPS13D^Halo and myc-Miro1 before and 2 hrs after Valinomycin addition. VPS13D, initially concentrated at mitochondria via its interaction with Miro, was shed from mitochondria, and relocated to the cytosol, upon treatment with Valinomycin, when EGFP-Parkin is recruited to mitochondria to initiate mitophagy. Scale bars= 30μm.

**Figure S3.**
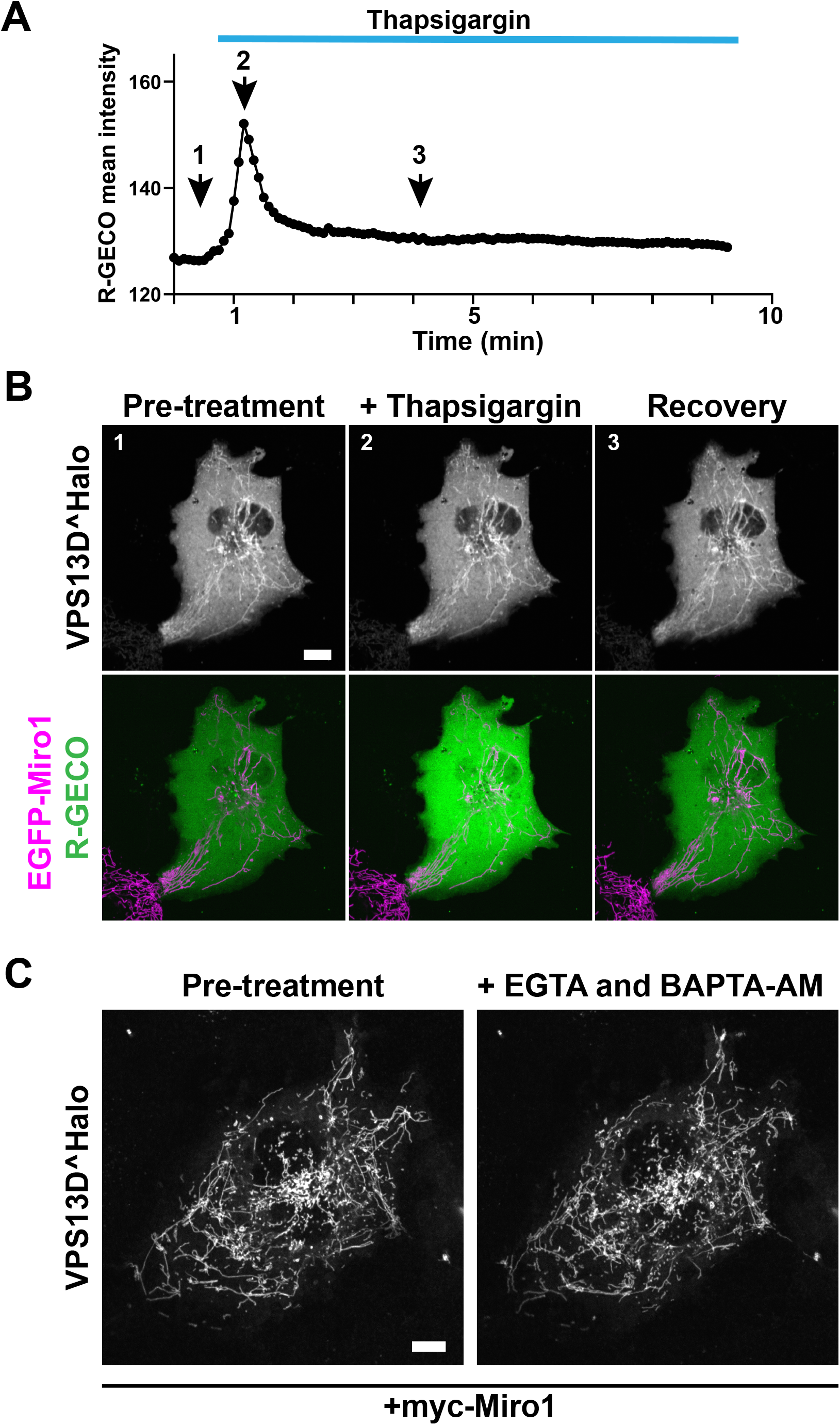
Recruitment of VPS13D to mitochondria by Miro is unaffected by changes in cytosolic Ca^2+^. (A and B) COS7 cell expressing EGFP-Miro1, VPS13D^Halo and the cytosolic Ca^2+^ sensor R-GECO. (A) Cytosolic Ca^2+^ levels before and after addition of the SERCA pump inhibitor thapsigargin. (B) Time-lapse confocal images showing snapshot from the timepoints indicated in (A) and demonstrating that the localization of VPS13D^Halo on mitochondria in the presence of overexpressed Miro1 is not affected by the Ca^2+^ concentration. (C) Time-lapse confocal images showing that the binding of VPS13D^Halo to mitochondria in the presence of co-expressed myc-Miro1 is unaffected by the addition of EGTA and BAPTA-AM to lower intracellular cytosolic Ca^2+^. Scale bars=10μm.

**Figure S4.**
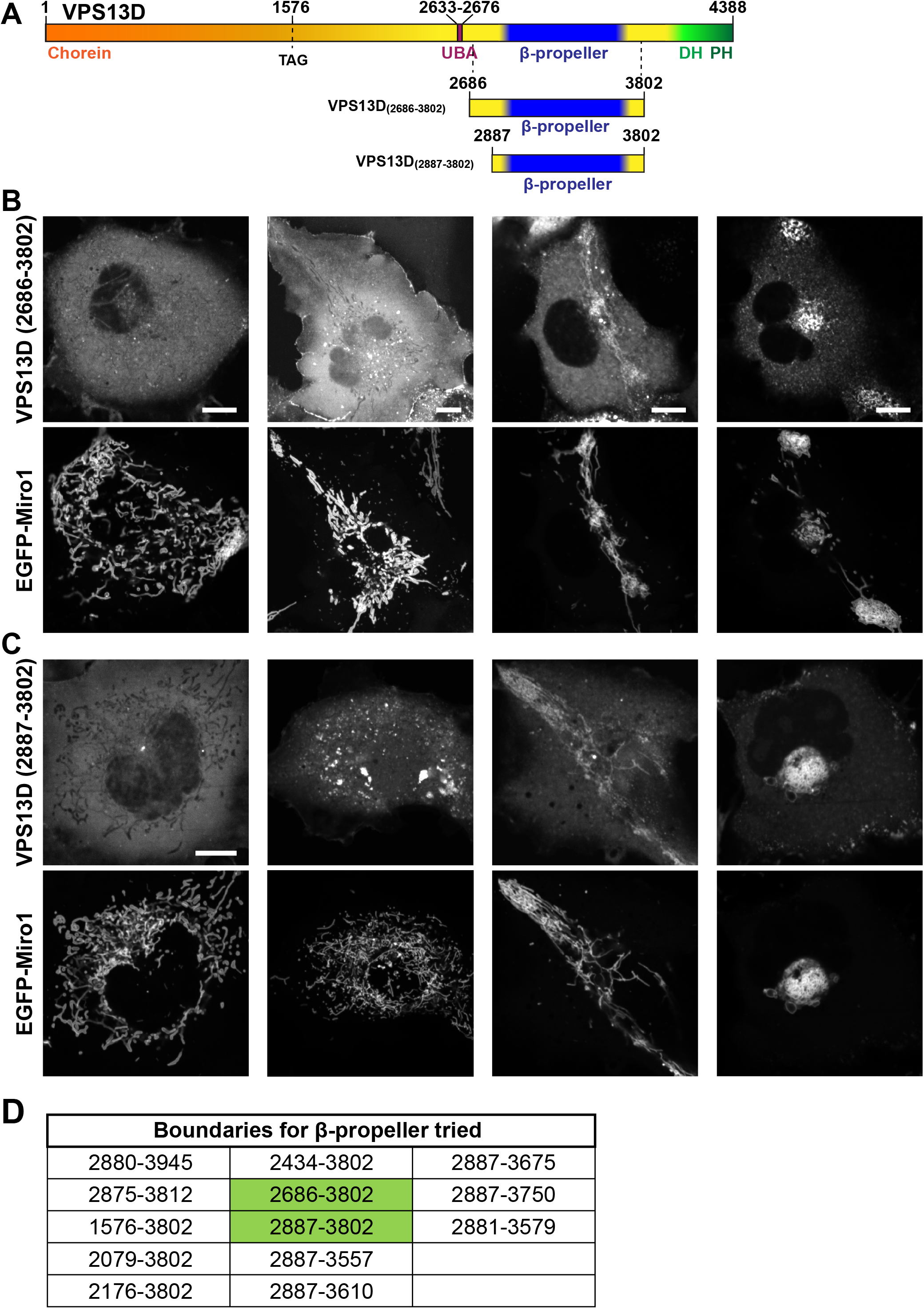
Localization of the constructs comprising the the β-propeller region of VPS13D only. (A) Cartoon depicting full length VPS13D and the VPS13D fragments used in the imaging experiment shown in B and C. (B-C) Confocal images of representative examples of the heterogenous localization of two mCherry-tagged VPS13D fragments in COS7 cells co-expressing EGFP-Miro1. The first column shows a cytosolic localization, the second column the presence of small aggregates and the third and fourth columns the colocalization of VPS13D with Miro on mitochondria whose localization is disrupted. Scale bars=10μm. (D) List of VPS13D fragments containing the β-propeller region that were tested for Miro-induced recruitment to mitochondria. All of them behaved similarly. The constructs used for field B and C are highlighted in green.

**Figure S5.**
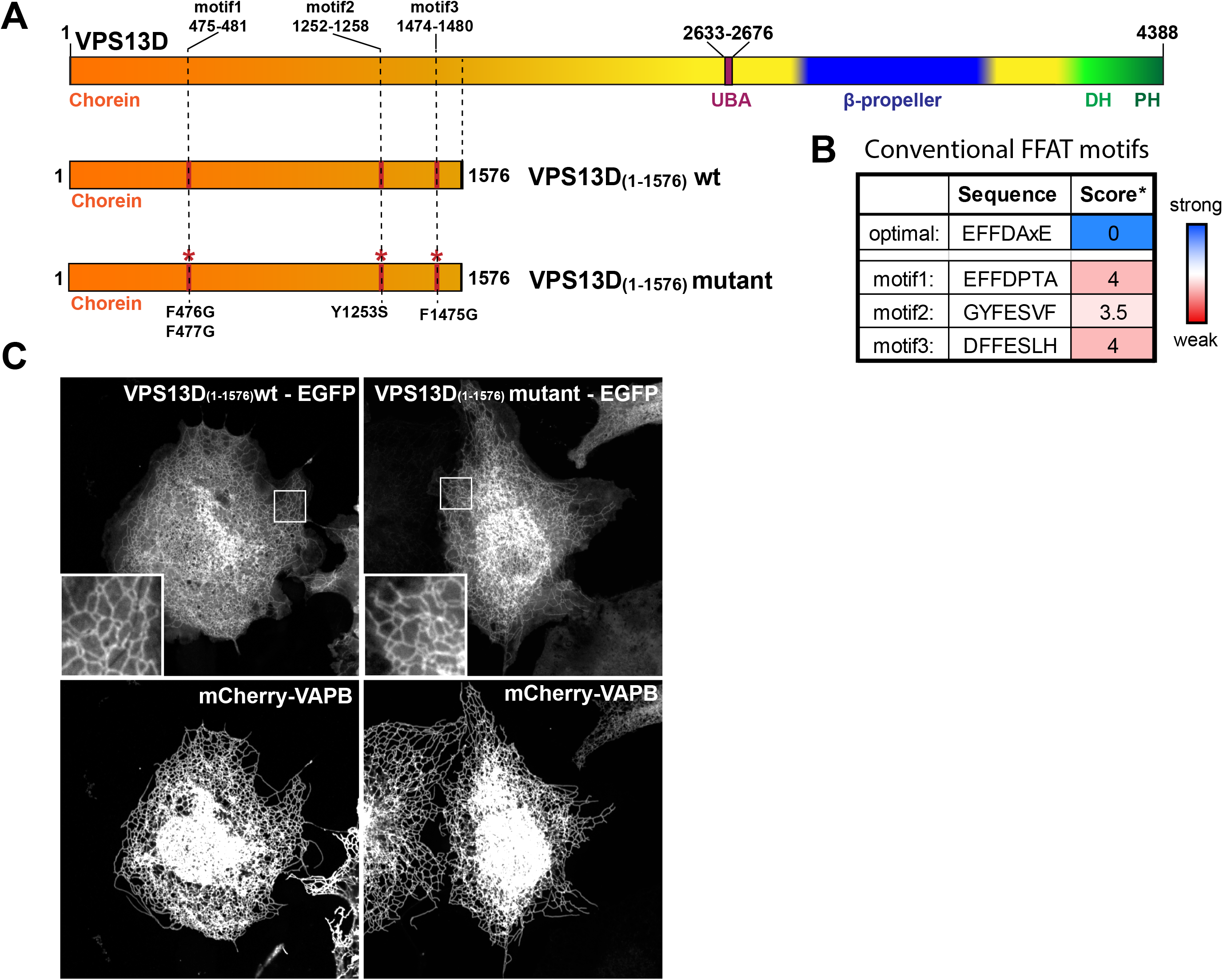
Predicted “conventional” FFAT motifs in the N-terminal region of VPS13D are not involved in the binding to VAP. (A) Cartoon depicting full length VPS13D and the constructs used for panel C, displaying the localization of the 3 predicted FFAT motifs and the mutations that were introduced to disrupt them. (B) Sequence and score of each of the 3 predicted conventional FFAT motifs.*Score was calculated using a previously described algorithm; scores 3.5 and 4 are considered weak FFAT motifs (Slee and Levine, 2019). (C) Co-expression of an EGFP-tagged N-terminal fragment of VPS13D (a.a. 1-1576) with the ER protein VAP-B shows robust recruitment of the fragment to the ER (left). The combined disruption of the 3 best predicted conventional FFAT motifs as indicated in panel (A) did not affect ER recruitment of this VPS13D fragment by VAP-B (right column). Scale bar=10μm.

## Materials and Methods

### DNA plasmids

A plasmid containing codon-optimized cDNA encoding human VPS13D, also including an mScarlet fluorescent protein after aminoacid residue 1576 flanked by BamHI restriction enzyme sites, was generated by and purchased from Genscript. This plasmid was linearized with BamHI and used to clone VPS13D^EGFP and VPS13D^Halo by Infusion cloning (Takara). An N-terminal fragment of VPS13D, Nterm(VPS13D)^EGFP, and a deletion mutant of VPS13D lacking the β-propeller region (VAB/WD40-like domain), VPS13D(Δβ-prop)^EGFP, were generated by PCR amplification and ligated into pEGFP-C1 by Infusion cloning, using EcoRI and AgeI sites. A C-terminally truncated VPS13D, VPS13D(ΔDHPH)^EGFP, was generated using site-directed mutagenesis (QuikChange II XL; Agilent technologies) by adding an early stop codon replacing the codon encoding aminoacid residue 3802. VPS13D constructs including the β-propeller region were generated by PCR amplification of the sequences encoding the different portions indicated in Fig S2B and ligation into a pmCherry-C1 backbone by Infusion cloning, using EcoRI and KpnI sites. Plasmids containing human Miro1(variant 1) and human Miro2 were a gift from P. Aspenström (Addgene #47888 and #47891). EGFP-Miro1, EGFP-Miro2 and mCherry-Miro1 were generated by PCR amplification of the coding sequence of Miro and ligated into pEGFP-C1(Addgene) or pmCherry-C1(Addgene), using EcoRI and XhoI sites. Halo-Miro1 was generated by PCR amplification of the coding sequence of Miro from EGFP-Miro1 and the coding sequence of the Halo protein from pHalo-C1 and ligated by Infusion cloning into a pcDNA3.1 backbone. For EGFP-Miro1v4 a double stranded DNA fragment encoding the aminoacid residues from exons 8 and 9 of Miro1 (i.e. the exons missing from Miro variant 1) was ordered from IDT and ligated into EGFP-Miro1 at residue 580 by Infusion cloning. For the optogenetic experiments, the cytosolic domain of Miro1 (Miro1(ΔTM)) and the iLID binding peptide (SspB, obtained from Addgene #60415, gift from B. Kuhlman) were fused into pmCherry-C1 using EcoRI and KpnI, by infusion cloning (mCh-Miro1(ΔTM)-SSPB). EGFP-Miro1 constructs containing mutations T18N, E208/E328K and S432N as well as VPS13D N-term fragments containing mutations of conventional FFAT motifs (F476/F477G, Y1253S, F1475G) or of the phospho FFAT motif (Y768A, T770A, T770D/P771A) were generated using site-directed mutagenesis (QuikChange II XL; Agilent technologies) of EGFP-Miro1. To generate BFP-VAP-B, the VAP-B coding sequence was amplified by PCR from a mCherry-VAP-B plasmid (our laboratory), the BFP coding sequence was amplified from mitoBFP (Addgene#49151) and were assembled by HiFi reaction (NEB). HALO-VAP-B was obtained by PCR amplification of VAP-B coding sequence from mCherry-VAP-B and pHALO-C1 using EcoRI and KpnI sites. Sialyltransferase(ST)-Halo was generated by PCR amplification of the ST coding sequence from ST-mRFP (our laboratory) and ligated into pHALO-N1 using EcoRI and KpnI sites. EGFP-Parkin was obtained by PCR amplification of rat Parkin cDNA (kind gift from E. Fon (McGill University, Montreal)) and ligated into pEGFP-C1 using EcoRI and XhoI. Other plasmids used in this study were kind gifts: mito-BFP from G. Voeltz (Addgene #49151), Venus-iLID-Mito from B. Kuhlman (Addgene #60413), mScarlet-SRL from D. Gadella (Addgene #85063). Mutant VAP-B (K87D, M89D) was previously generated in our lab(Dong et al., 2016).

### Antibodies and reagents

Primary antibodies were obtained as follows: anti-RHOT1 (H00055288-M01; Abnova), anti-Miro2 (ab224089; abcam), anti-TOMM40 (18409-1-AP; Proteintech), anti-VPS13D (ab202285; abcam), anti-GAPDH (40-1246; Proteus Biosciences Inc); anti-GFP, (ab290; abcam), anti-myc (sc-40; Santa Cruz Biotechnology), anti-RFP (600-401-379; Rockland Inc). Halo tag ligands were a kind gift from L. Lavis (Janelia Research Campus, Ashburn, VA). Valinomycin was purchased from Sigma-Aldrich and used at 10μM concentration. RNAis for Miro1 were purchased from Ambion (#4390824) and the ones for Miro2 from IDT (hs.Ri.RHOT2.13). Specific primers were purchased from IDT, for sequences refer to Supplementary Table 1.

### Cell culture and transfection

COS7 and HeLa cells (from ATCC) were cultured at 37°C and 5% CO2 in DMEM containing 10%FBS, 1mM sodium pyruvate, 100U/ml penicillin, 100mg/mL streptomycin and 2mM L-glutamine (all from Gibco). For imaging experiments, cells were seeded on glass-bottomed dishes (MatTek) at a concentration of 75×10^3^ cells per dish and transiently transfected after 6h using FuGene HD (Promega). For Miro1 and Miro2 knockdown experiments, 12h following the transient transfection with VPS13D and mitoBFP, HeLa and COS7 cells were treated RNAi (30pmol/gene) using Lipofectamine RNAiMAX (Life Technologies) and imaged 36h after.

### Neuronal cultures

Experiments were performed in accordance with the Yale University Institutional Animal Care and Use Committee. Briefly, hippocampi were dissected from P0 mouse brains, neurons were dissociated by papain treatment and seeded in serum-based medium on poly-D-lysine coated, glass-bottomed dishes (MatTek), as previously described {Sun 2019}. After six hours, the serum-based medium was removed and replaced with neuronal growth media [Neurobasal A supplemented with B27 and Glutamax (all from Gibco)]. After 11 DIV, cells were transfected using Lipofectamine 2000 (Life Technologies).

### Microscopy

#### Live cell imaging

Just before imaging, the growth medium was removed and replaced with Live Cell Imaging solution (Life technologies). All live cell imaging was carried out at 37°C and 5%CO_2_. Spinning-disk confocal microscopy was performed using an Andor Dragonfly system equipped with a PlanApo objective (63X, 1.4NA, Oil) and a Zyla sCMOS camera. For the hypotonic lysis experiments, Live Cell Imaging solution was replaced with distilled water and cells were imaged at a rate of 0.5Hz.

For experiments involving cytosolic Ca^2+^ changes, COS7 cells were seeded and transfected as explained above. To acutely increase cytosolic Ca^2+^, 2μM thapsigargin (Life Technologies) was added to the Live Cell Imaging solution (Life technologies), which contains 1.8 mM Ca^2+^) and cytosolic Ca^2+^ was monitored by the intensity of the sensor R-GECO (plasmid was a gift from R. Campbell, Addgene #45494). To decrease cytosolic Ca^2+^, 4mM EGTA and 10μM BAPTA-AM (Thermo Fisher) was added to the medium during imaging. Calcium experiments were repeated twice, in at least 4 cells for each case.

#### Optogenetic experiments

For whole cell activation experiments, an Andor Dragonfly system (see above) was used and recruitment to mitochondria was achieved with a single 200ms pulse of the 488nm laser. For localized activation to promote recruitment to a single mitochondrion, an Improvision UltraVIEW VoX system (Perkin Elmer), built around a Nikon Ti-E inverted microscope and controlled by Volocity software (Improvision), was used. Imaging was carried out at 37°C with a 63x PlanApo oil objective (1.45 NA). A built-in photo-perturbation unit was used to deliver 488nm light pulses in a 5μm^2^ area.

### Image processing, analysis and statistics

Fluorescence images were processed using FIJI (ImageJ; NIH) software. *Gaussian Blur* filters were applied on some of the images presented. The fluorescent signal in the Supplementary Movies were corrected for photobleaching using the *Bleach Correction* function in FIJI.

For the quantification of the optogenetic experiment, kymographs were built tracing a line across the mitochondria in the unprocessed images, resulting in images like the examples shown in Fig 1G, from which the intensity profile was measured. The fluorescence intensities were normalized to the min and max values using the following formula: *Normalized* 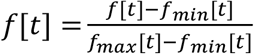, where *f*[*t*] = *Fmito*[*t*] − *Fbkgd*[*t*].

For the analysis of VPS13D mitochondrial recruitment, cells co-expressing VPS13D^EGFP and mitoBFP were imaged and analyzed individually, using an automated macro designed in FIJI. *Gaussian Blur* and *Enhance Local Contras*t were applied to the mitoBFP channel to generate an accurate mask of mitochondria. The total amount of VPS13D signal on mitochondria was then obtained by measuring the VPS13D^EGFP intensity within that mask. To obtain a mask covering the cytosolic area surrounding mitochondria the *Dilate* function was applied to the mitochondrial mask, specifically, a 2-times dilated mitochondrial mask was subtracted from a 3-times dilated one. The resulting mask, a 1-pixel wide mask covering the area lining the profile of mitochondria, was then used to measure VPS13D^EGFP intensity in the cytosol surrounding mitochondria. This second value was used to normalize for expression levels. The ratio between the intensity of VPS13D on mitochondria and the intensity of VPS13D in the cytosol surrounding them was plotted on graph as the value of VPS13D enrichment at mitochondria. For statistical analysis GraphPad Prism 8 software was used. Brown-Frosythe and Welch’s ANOVA test was used to assess significant differences among experimental groups, followed by a Games-Howell’s multiple comparisons test.

## Movie legends

Movie S1: Confocal time-lapse imaging of a COS7 cell co-expressing VPS13D^Halo, mCh-Miro1(ΔTM)-SspB and EGFP-iLID-Mito (not shown) showing recruitment of VPS13D and Miro1 upon blue light irradiation (at 0s) of the area indicated by the white box. Scale bars=10μm.

Movie S2: Confocal time-lapse imaging of a COS7 cell co-expressing VPS13D^Halo, EGFP-Miro1 and BFP-VAP-B upon hypotonic shock. Mitochondria and ER vesiculate but remain tethered by hot spots of VPS13D. Cells were imaged right after water addition at a rate of 0.5 fps for a total of 2 minutes. Some frames were lost due to focal plane changes. Scale bar=3μm.

## Acknowledgements

We thank Y. Wu, M. Hammarlund, H. Falahati, and J.H. Park for discussion. We also thank F. Wilson and A. Dao for outstanding technical assistance. This work was supported in part by NIH grant NS36251, the Kavli Foundation, the Parkinson’s Foundation (PF-RCE-1946) and MJFF (ASAP-000580) to P.D.C and by a grant from the CZI (#2020-221912) to P.D.C and H.S.. A.G.S. was supported in part by a Jung-Stiftung Scholarship.

---

**Table S1:**
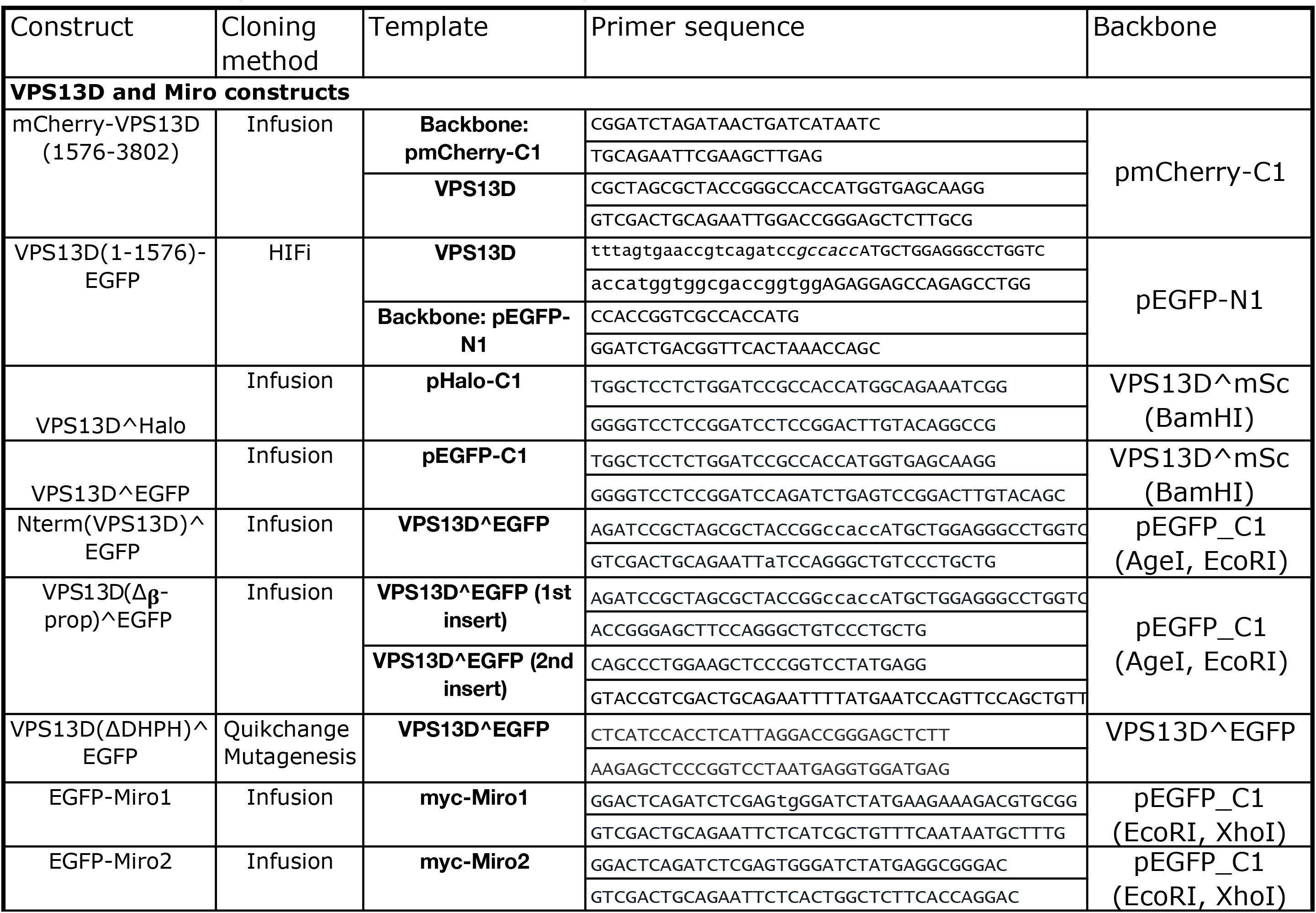
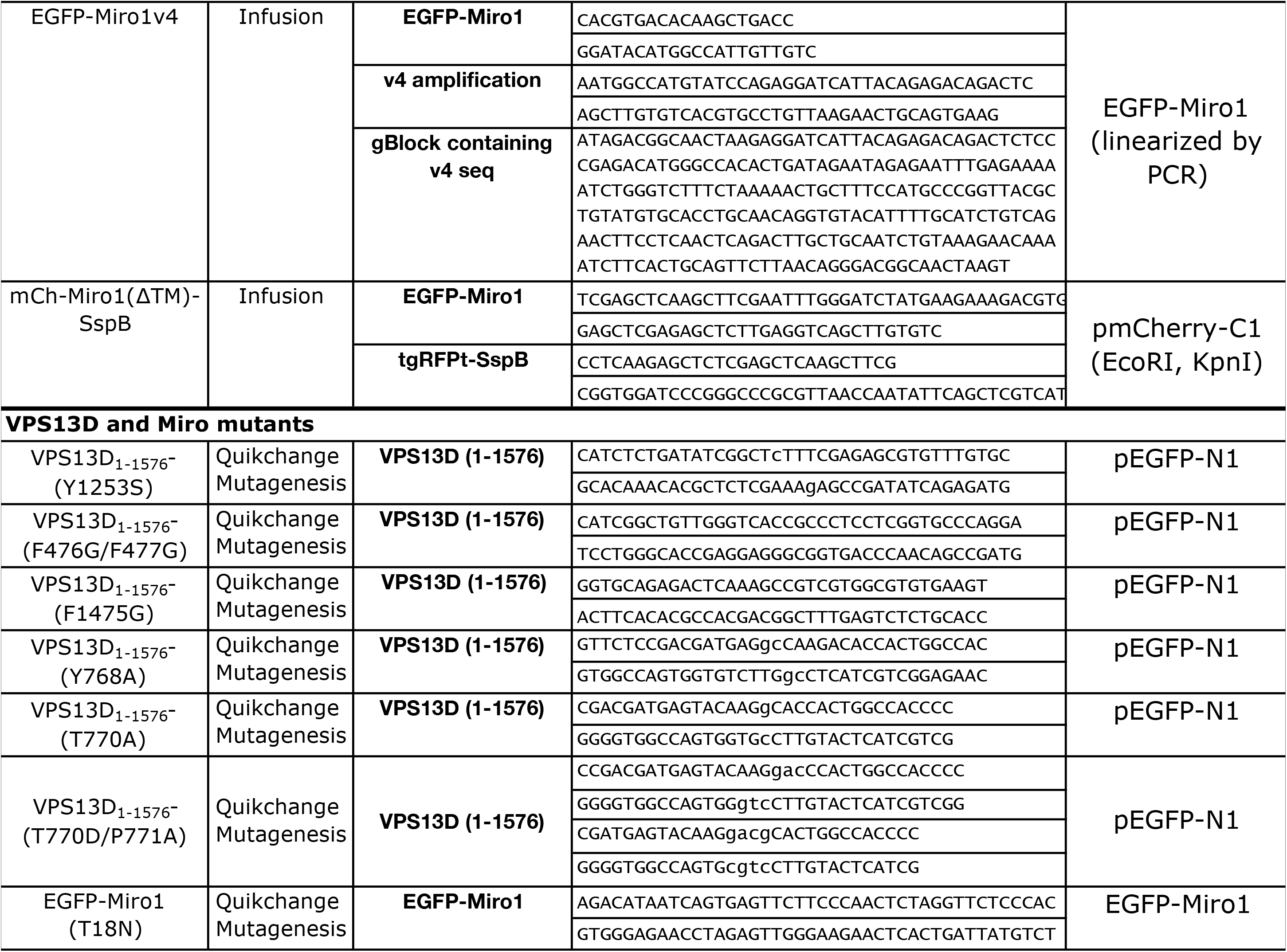
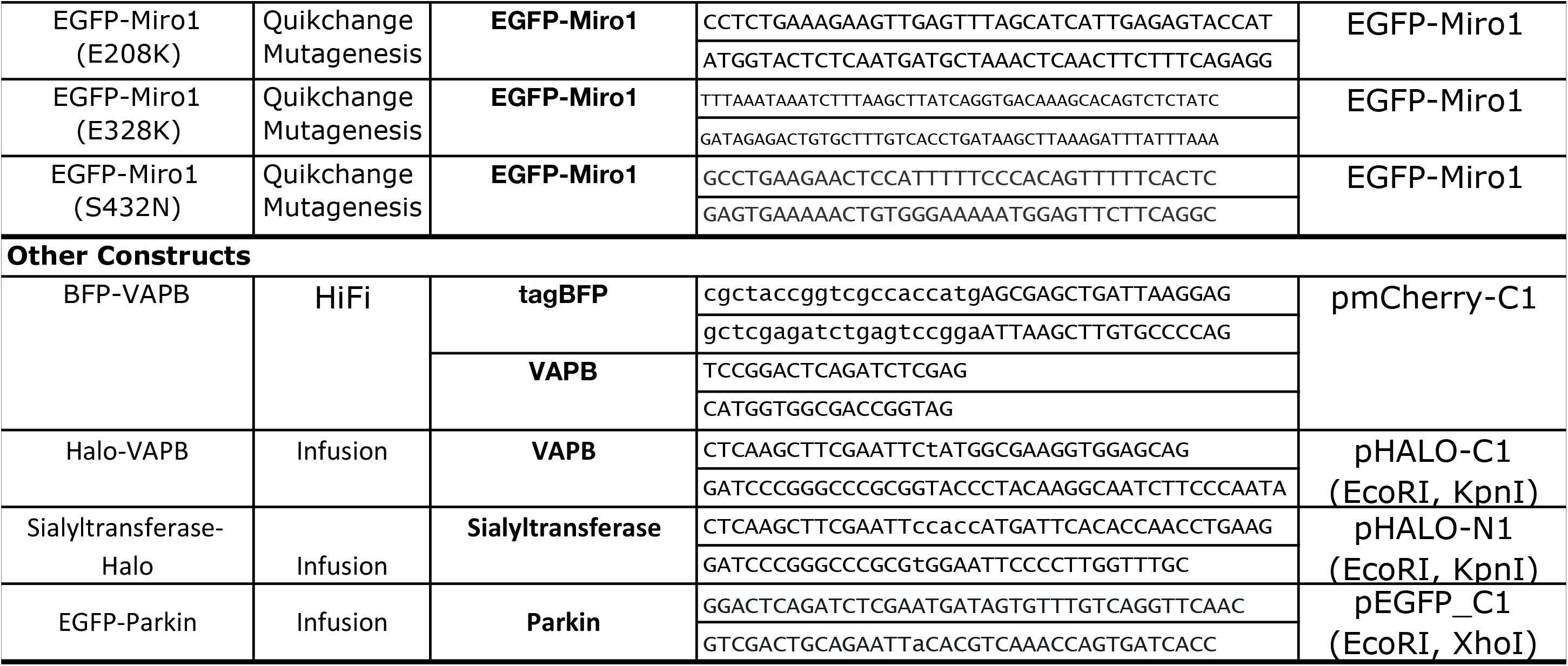
List of primers used in this study

## Notes

### Competing Interest Statement

The authors have declared no competing interest.

